# Comparison of Two-Talker Attention Decoding from EEG with Nonlinear Neural Networks and Linear Methods

**DOI:** 10.1101/504522

**Authors:** Gregory Ciccarelli, Michael Nolan, Joseph Perricone, Paul Calamia, Stephanie Haro, James O’Sullivan, Nima Mesgarani, Thomas Quatieri, Christopher J. Smalt

## Abstract

Auditory attention decoding (AAD) through a brain-computer interface has had a flowering of developments since it was first introduced by Mesgarani and Chang (2012) using electrocorticograph recordings. AAD has been pursued for its potential application to hearing-aid design in which an attention-guided algorithm selects, from multiple competing acoustic sources, which should be enhanced for the listener and which should be suppressed. Traditionally, researchers have separated the AAD problem into two stages: reconstruction of a representation of the attended audio from neural signals, followed by determining the similarity between the candidate audio streams and the reconstruction. In this work, we compare the traditional two-stage approach with a novel neural-network architecture that subsumes the explicit similarity step. We compare this new architecture against linear and non-linear (neural-network) baselines using both wet and dry electroencephalogram (EEG) systems. Our results indicate that the wet and dry systems can deliver comparable results despite the latter having one third as many EEG channels as the former, and that the new architecture outperforms the baseline stimulus-reconstruction methods for both EEG modalities. The 14-subject, wet-electrode AAD dataset for two competing, co-located talkers, the 11-subject, dry-electrode AAD dataset, and our software are available to download for further validation, experimentation, and modification.

## 1 Introduction

Hearing loss, and the associated use of hearing-aids, is rising among the general population [1], and as shown by recent statistics from the US Dept. of Veterans Affairs, is particularly prevalent among retired military personnel [2]. Despite widespread use of hearing aids, and the incorporation of spatial and spectral algorithms for noise reduction, hearing-aids often are considered unsatisfactory in regard to their performance in noisy environments [3–5]. Particularly when background noise includes other talkers, hearing aids suffer because they have difficulty separating the “signal” (*i.e*., the talker of interest to the listener) from the “noise” (*i.e*., all other talkers) due to similarities in spectro-temporal characteristics. The failure of hearing aids to improve listening ability in complex acoustic environments, either due to poor device performance, or lack of use triggered by poor performance, is associated with social isolation and various forms of cognitive decline such as depression [6–8]. Therefore, solving the problem of assisted listening in multi-talker environments could have wide societal benefits in terms of communication and mental health. Auditory attention decoding (AAD) is a recent approach aimed at such a solution, one which exploits knowledge of the listener’s auditory intent (attention) to isolate and enhance the desired audio stream and suppress others.

Evidence for neural encoding of speech has been shown with various sensing modalities including electroencephalography (EEG) [9], magnetoencephalography (MEG) [10], and electrocorticography (ECoG) [11]. The exploitation of such encoding for AAD in a two-talker paradigm was initially demonstrated by Mesgarani and Chang [12], through a classifier acting on speech spectrograms reconstructed from ECoG data. Comparison of the predicted spectrograms with those from the actual speech sources provided the identity of the attended talker with 93% accuracy when the subjects were known to be attending to the instructed stimulus. Since then, AAD has been achieved successfully with many variations on this initial technique.

The most common approach to AAD, first described in [13] and depicted in Fig 1, involves EEG for capturing neural data as a more practical and less invasive modality than ECoG. The approach uses a linear least-squares method for stimulus (broadband speech envelope) reconstruction and correlation of actual and predicted speech envelopes to identify the attended talker. Stimulus reconstruction is also known as the “backward” problem in AAD, as the mapping from EEG to stimulus is the reverse of the natural auditory stimulus/response phenomenon. By contrast, predicting EEG from the stimulus is known as the “forward” problem.

**Fig 1.**
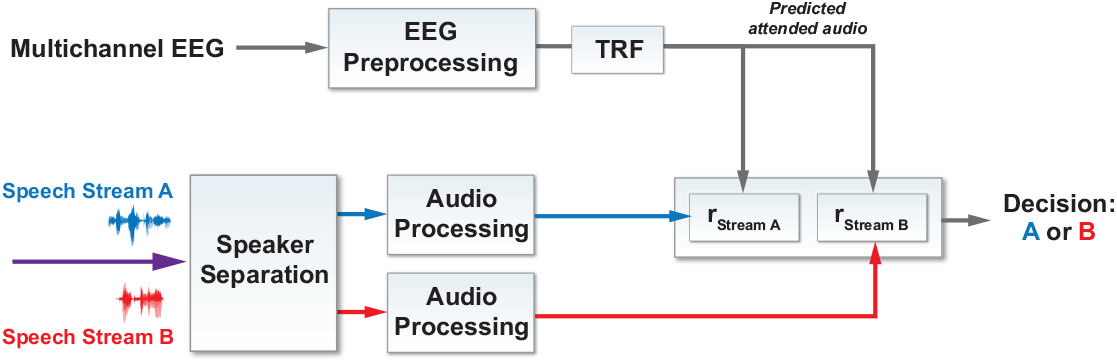
System architectures for auditory attention decoding: backward model. The temporal response function (TRF) can be linear (least squares, see Eq. 1) or non-linear (neural network, see Fig 3).

The attention decision typically is between two simultaneous, spatially separated talkers. This approach has been modified to evaluate: sensitivity to number of EEG channels and size of training data [14]; robustness to noisy reference stimuli [15, 16]; the use of auditory-inspired stimulus pre-processing including subband envelopes with amplitude compression [17]; cepstral processing of EEG and speech signals for improved correlations [18]; the effects of speaker (spatial) separation and additional speech-like background noise [19]; the effects of (simulated) reverberation [20]; and potential performance improvements through various regularization methods [21].

Considering the AAD pipeline as comprising steps for neural data acquisition, stimulus representation, signal processing (*e.g*., forward or backward predictive modeling), and attention determination, alternate techniques have been described with variations of each of these components. MEG [22] and ECoG [23] continue to serve as neural sensing modalities, while EEG channels have been reduced in number in an effort to move toward less obtrusive, portable systems [24, 25]. Speech stimuli have been represented with spectrograms [23] and frequency-dependent envelopes after gammatone filtering [26]. To exploit the power and biological relevance of non-linear processing, effective implementations of the backward model with neural networks have been shown [27], and while much less popular, linear versions of the forward model (predicting EEG from the stimuli) are described in [21, 25]. As an alternative to both forward and backward modeling, canonical correlation analysis, which involves transforming both stimulus and response to maximize mutual projections and thus improve correlations, has been applied to EEG and audio data, both with various filters, to enhance AAD performance [28]. Finally, state-space models have been applied as a final step in AAD systems to smooth noisy attention decisions and allow for near real-time update rates [29].

Measuring the performance of AAD systems typically involves an intuitive computation of decoding accuracy, *i.e*., the percentage of decoding opportunities for which the system correctly identifies the attended talker. Overall results often are generated with a leave-one-out cross-validation scheme iterated over the collected dataset. This approach is used in both the backward [13–15, 17, 21] and forward [25] modeling paradigms. System accuracy also has been reported for predicting the *unattended* talker [13, 20], but in both cases performance is worse than that for predicting the attended talker. In [29], the *[lscript]*_1_-norm of the attended and unattended decoder coefficients are used as “attention markers” to generate a smooth, near real-time (∼2-second latency) attentional probability through a state-space estimator. Talker classification is considered correct if the probability estimate *and* its 90% confidence interval for the attended talker are above 0.5, and accuracy is again measured as the percentage of correctly classified opportunities. In [21, 27], performance is reported as an information transfer rate, *i.e*., the number of correct decoding decisions per minute.

Comparison of performance statistics across different published results, even those using the same decoding approach and performance metric, is hampered by variations in experimental parameters including talker number, angular separation, and gender, as well as number/placement of EEG electrodes, and by variations in processing parameters such as EEG or speech-envelope bandwidths, and correlation lags and window sizes. To address these barriers, in this paper we describe two datasets and three decoding algorithms along with results from each of the six combinations. The datasets include wet and dry EEG data collected from 14 and 11 subjects, respectively, during an auditory-attention experiment with two simultaneous, co-located talkers (one female, one male). The algorithms include a linear least-squares stimulus-reconstruction decoder described in [13], a neural-network stimulus-reconstruction decoder described in [27], and a novel convolutional neural-network classifier that predicts the attended talker without explicit forward or backward prediction (Fig 2).

**Fig 2.**
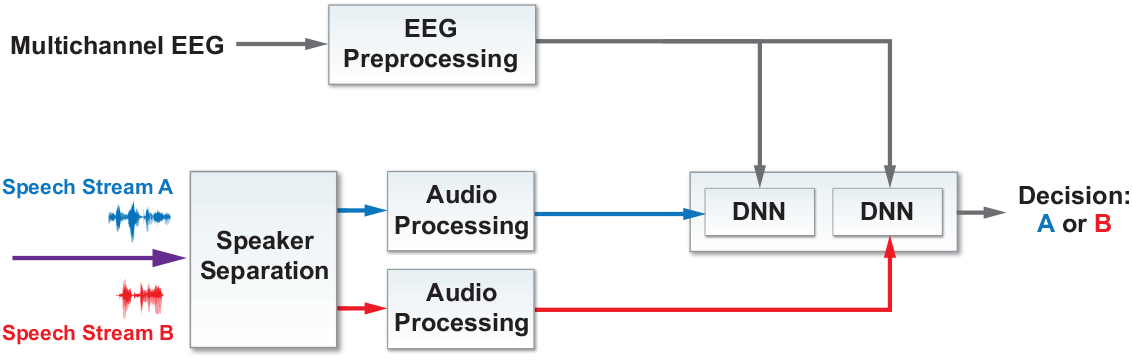
System architectures for auditory attention decoding: DNN binary classification. See Fig 4 for a specific instance of the DNN.

## 2 Methods

### 2.1 AAD Experimental Collection

#### 2.1.1 Protocol

Speech from two co-located talkers, one male, and one female, was presented to each subject in a quiet, electrically shielded audiometric booth. The audio was presented from a single loudspeaker directly in front of the subject, with the stimuli lasting approximately 30 minutes. The stimuli consisted of four “wikiHow.com” instructions lasting approximately 5 minutes each: “How to Make Waffles”, “How to Take Care of a Dog”, “How to be a Shepherd”, and “How to Identify Birds”. Each story (attended audio) was heard twice, once read by the male and once by the female talker, with a different story by the opposite gender presented simultaneously as the distractor (unattended) audio stream. The order of the two talkers, as well as the attended and distractor audio streams were randomized for each subject. Participants were instructed as to which gender talker to focus on at the start of each story on a screen in front of them throughout the experiment. Each story was interrupted randomly after 5–10 sentences were presented, and the participant was asked to repeat the last sentence of the attended talker. We term each uninterrupted listening interval as a “part”. A subset of subjects also participated in an auditory oddball task, but that data is not part of this analysis.

#### 2.1.2 Subjects

Fourteen MIT Lincoln Laboratory employees (9 male, 5 female) gave written, informed consent to participate in the experiment, in a protocol approved by the MIT Committee on the Use of Humans as Experimental Subjects and the US Army Medical Research and Materiel Command (USAMRMC) Human Research Protection Office. Eleven subjects, partially overlapping with the original fourteen, agreed to participate in a second experiment with the same protocol as the first. The first experiment used a wet EEG system, and the second used a dry EEG system (see Section 2.1.3 below). Most participants self-reported normal hearing; two subjects reported known hearing loss.

To ensure that subjects were on task, as well as potentially to exclude subjects that were unwilling or unable to attend to the target speaker, we checked the randomized interruptions of the stimuli presentations for a qualitative measure of attention. No subjects were excluded due to performance concerns.

#### 2.1.3 EEG Instrumentation and Preprocessing

Wet electrode EEG data were collected using a Neuroscan 64-channel Quik-Cap and a SynAmps RT amplifier with a sampling rate of 1000 Hz, and recorded in Curry data-acquisition software (Compumedics, Charlotte, NC). Additional electrodes were placed on both mastoids, as well as above, below, and next to the left eye. The reference electrode was located halfway between CZ and CPZ. Dry electrode EEG data were collected using a Wearable Sensing DSI-24 system (San Diego, CA), a joint sensor platform and signal amplifier. The system records from 18 scalp channels and two reference channels attached to the subject’s earlobes. Data were collected at a 300 Hz sampling rate using DSI-Streamer software.

Prior to analysis, all EEG data were down-sampled to 100 Hz using MATLAB’s resample function (Mathworks, Natick, MA), which applies an anti-aliasing low-pass filter with a cutoff frequency of 50 Hz. EEG data were band-pass filtered with a passband frequency of 2 to 32 Hz.

#### 2.1.4 Audio Preprocessing

For both the stimulus reconstruction and binary classification methods, we pre-processed the two clean, audio streams to extract their broadband envelopes using the iterative algorithm in [30]. Envelopes were subsequently downsampled to a 100-Hz sampling rate.

### 2.2 Linear Decoding

To recreate the linear, stimulus-reconstruction approach in [13] (see Fig 1), we implemented a regularized, least-squares transform (LSQ) from EEG response data to audio envelope according to the ridge regression equation: 

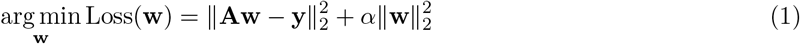

EEG data segments are stacked in rows of the **A** matrix. Each row vector contains all the time points of the context window for all the EEG channels. **y** is a column vector of the audio envelope. Each EEG row vector is transformed or decoded by a weight vector **w** into the audio envelope sample that corresponds to the most recent time sample in the row vector.

The LSQ weights, **w**, are often called the temporal response function (TRF) from the response-prediction EEG literature in which the EEG is seen as a response to the audio stimulus. Strictly speaking, when attention decoding is formulated in the backwards direction, the weights represent an inverse TRF.

The regularization parameter,*α*, was selected on a per-subject, per-test-part basis from a set of three heuristically chosen values. A robust standard scaling was applied to the training and testing audio and EEG data, also on a per-subject, per-test-part basis, using the estimated median and inter-quartile range of the training data. Each segment of data used for the LSQ method (and the DNN correlation-based method) was 26 samples long (approximately 250 ms given the 100-Hz sampling rate). Estimation was performed using Scikit-learn’s `linear_model.RidgeCV` method [31]. Separate models were trained for each subject; no transfer learning across subjects was used in this analysis.

### 2.3 Nonlinear Decoding

The motivation for applying a deep neural network (DNN) to the AAD problem is that a non-linear decoder may provide improved performance relative to a linear decoder due to the inherent non-linear processing of acoustic signals along the auditory pathway. A DNN is a prototypical non-linear method flexible enough to handle multi-dimensional time series data. We use a neural network inspired by [27] for the correlation-based classifier, and a novel convolutional DNN for the integrated classification decision architecture.

#### 2.3.1 Neural Network for Stimulus Reconstruction

A simple neural-network architecture comprising a single hidden layer with two nodes was shown in [27] to yield the best performance from a group of more complicated networks considered. Our adaptation of that network, shown in Fig 3, includes batch normalization [32] before the inputs to each layer, and a hard hyperbolic tangent (as opposed to a linear function) for the output layer’s activation to enforce our prior expectation that the audio envelope be bounded.

**Fig 3.**
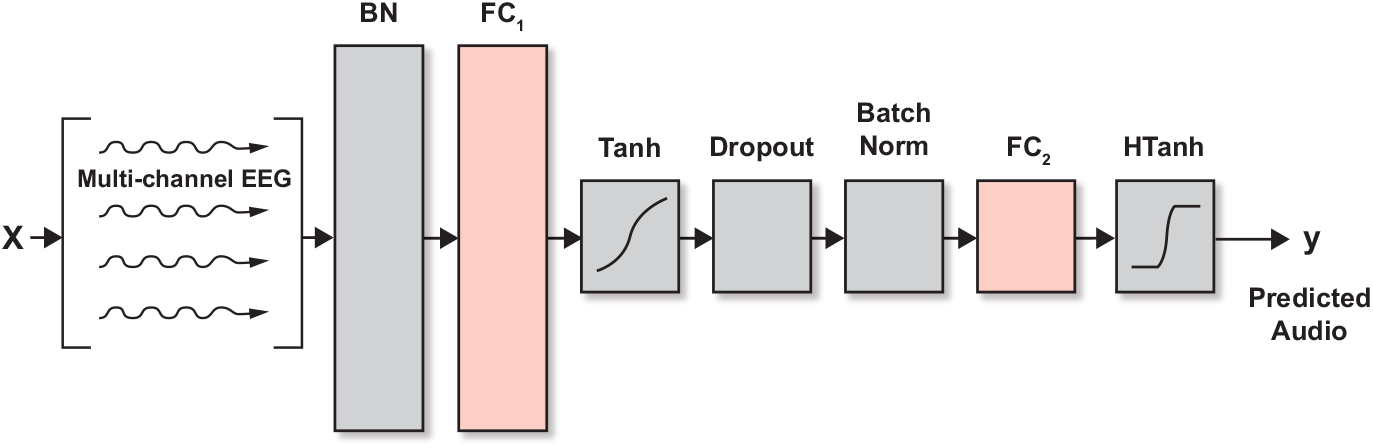
The neural network architecture for stimulus reconstruction, based on the design in [27]. There is one hidden layer with two nodes (FC_1_) to enforce significant compression of EEG data before being transformed to a predicted audio stimulus (see Fig 1 for the system architecture). BN = batch normalization, FC = fully connected.

The network was trained with the Adam optimizer using a batch size of 8192*8 samples, weight decay of 10, a learning rate of 10^−3^ for the first 150 steps, and then a learning rate of 10^−4^ for the remaining steps for a total of 250 steps. These parameters were heuristically chosen by inspecting intermediate train and validation-set loss curves where two additional parts were reserved from within the train set for validation. Following [27] we also employed a correlation-based loss function rather than a mean-squared error-loss function to exploit the prior knowledge that we ultimately will be testing the reconstructed waveform and AAD performance with a correlation metric.

#### 2.3.2 Neural Network for Direct Classification

Our novel end-to-end classification network with integrated similarity computation between EEG signals and a candidate audio envelope is pictured in Fig 4. It comprises two convolutional layers, the first of which uses a kernel of three samples, and the second of which uses a kernel of one sample. The convolutional layers are followed by a set of four, fully connected layers that decrease in size in the later stages. We use batch normalization and dropout [33] throughout, and the exponential linear unit [34] for the non-linearity. Training includes a binary cross-entropy loss function, batch size of 1024, Adam optimizer, no weight decay, and a learning rate of 10^−3^. We terminated the optimization process if the loss on the training set declined to below 0.09 or if the optimizer had run for 2400 steps. Because of computational limits on our computers, we randomly downsampled the 10-second set of samples over which a frame was evaluated by a factor of four.

**Fig 4.**
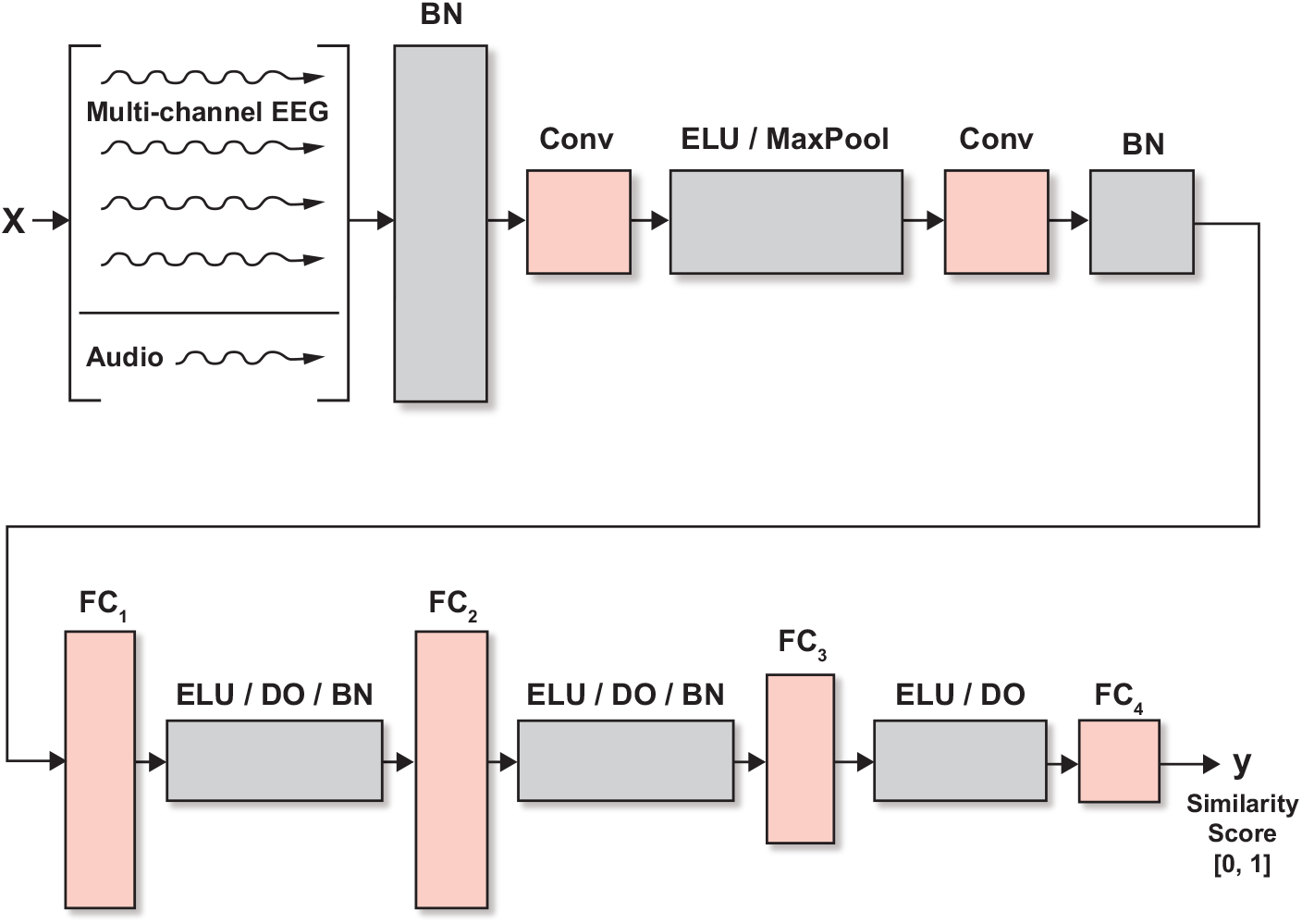
The convolutional architecture used for integrated similarity computation between EEG and a candidate audio stream. Components include batch normalization (BN), convolution layers (Conv_*i*_), exponential linear units (ELU), drop-outs (DO), and fully connected layers (FC_*i*_). Wet EEG (kernel, num ch in, num ch out): Conv_1_: 3×65×64, Conv_2_: 1×64×2, Dry EEG: Conv_1_: 3×19×19, Conv_2_: 1×19×2, Both: FC_1_: 246×200, FC_2_: 200×200, FC_3_: 200×100, FC_4_:100×1, MaxPool 1D, stride:2. See Fig 2 for the system architecture.

### 2.4 Methods of Evaluation

#### 2.4.1 Correlation-Based Evaluation

Algorithm performance was evaluated in a leave-one-out cross-validation paradigm across all audio parts presented to the subject. Multi-part training was performed by concatenating the presented audio data and recorded EEG response data. The concatenation was performed after each part was converted into a data matrix for the algorithm estimation to avoid discontinuities. The LSQ (linear) and DNN (non-linear) estimators were trained to reconstruct the attended audio using the training audio and EEG. Then, given the test EEG, each algorithm attempted to reconstruct the attended audio stimulus.

The estimated audio was then compared to the two candidate audio streams (attended and unattended) using Pearson correlation. The correlation was computed for ten-second, non-overlapping windows for the test part. If the left-out part was less than ten seconds, it was not evaluated. Decoding accuracy was computed as the percentage of 10-second windows for which the correlation coefficient with the attended audio envelope was higher than the correlation coefficient with the unattended audio envelope.

#### 2.4.2 Classification-Based Evaluation

In the DNN classification architecture, the algorithm directly makes a similarity prediction between the recorded EEG and each of the candidate audio streams. In other words, the similarity metric is learned by the network during the training rather than dictated by the user. Given the similarity scores for each candidate audio stream, the attended stream is declared as the one with the highest score. To keep the decision rate the same between the two network architectures, we provide the classification algorithm data segments that are ten seconds in duration.

## 3 Results

### 3.1 Decoding Accuracy

Decoding results for the wet EEG system are shown in Fig 5, and for the dry EEG in Fig 6. Each figure shows the per-subject average decoding accuracy using the linear correlation, neural-network based correlation, and DNN classification methods. Chance-level performance, indicated by the black stars, was computed as the 95^*th*^ percentile point of a binomial distribution with *p* = 0.5 and *n* equal to the number of non-overlapping 10-second windows. Mean decoding accuracies across subjects are summarized in Table 1. A 2-way mixed-model ANOVA (EEG Type by Algorithm Type) was performed with subjects modeled as a random factor. We found a main effect for the choice of algorithm type (*F* (2, 56) = 73.5*, p* < 0.0001) but not for EEG type (*F* (1, 56) = 0.02*, p* = 0.89). The interaction between algorithm choice and EEG type was also significant (*F* (2, 56) = 5.8*, p* < 0.01). Bonferroni corrections were used for *post-hoc* multiple comparisons, and revealed statistically significant differences between the DNN classifier and both stimulus-reconstruction algorithms for both wet and dry EEG. There was no significant pairwise effect of the EEG type for any of three algorithms tested.

**Fig 5.**
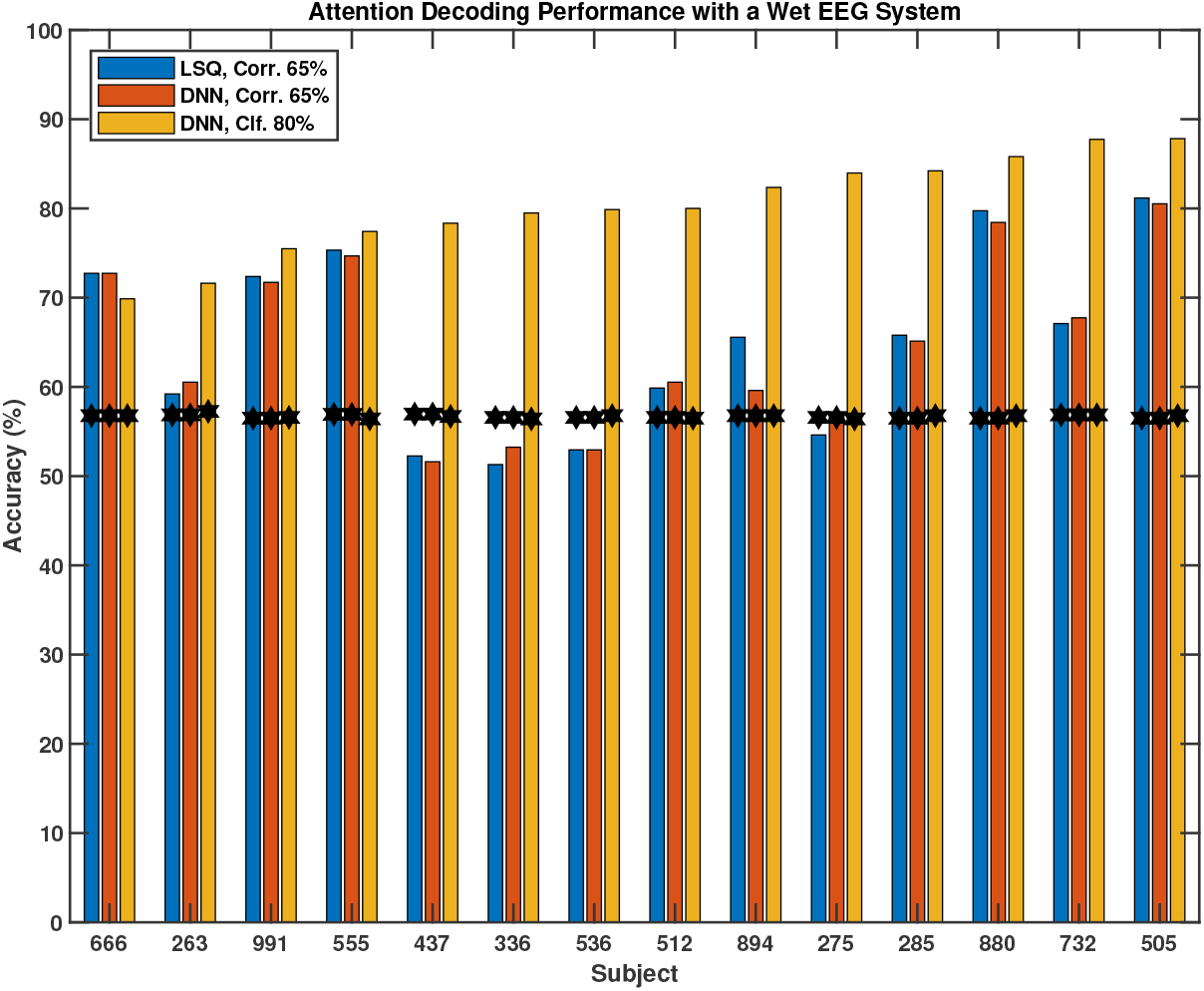
Per-subject attention-decoding accuracy using a wet EEG system. 10-second evaluation window, three algorithms: linear stimulus reconstruction (LSQ Corr.), non-linear stimulus reconstruction (DNN Corr.), and DNN classification (DNN Clf.). Chance performance is indicated by the black stars.

**Fig 6.**
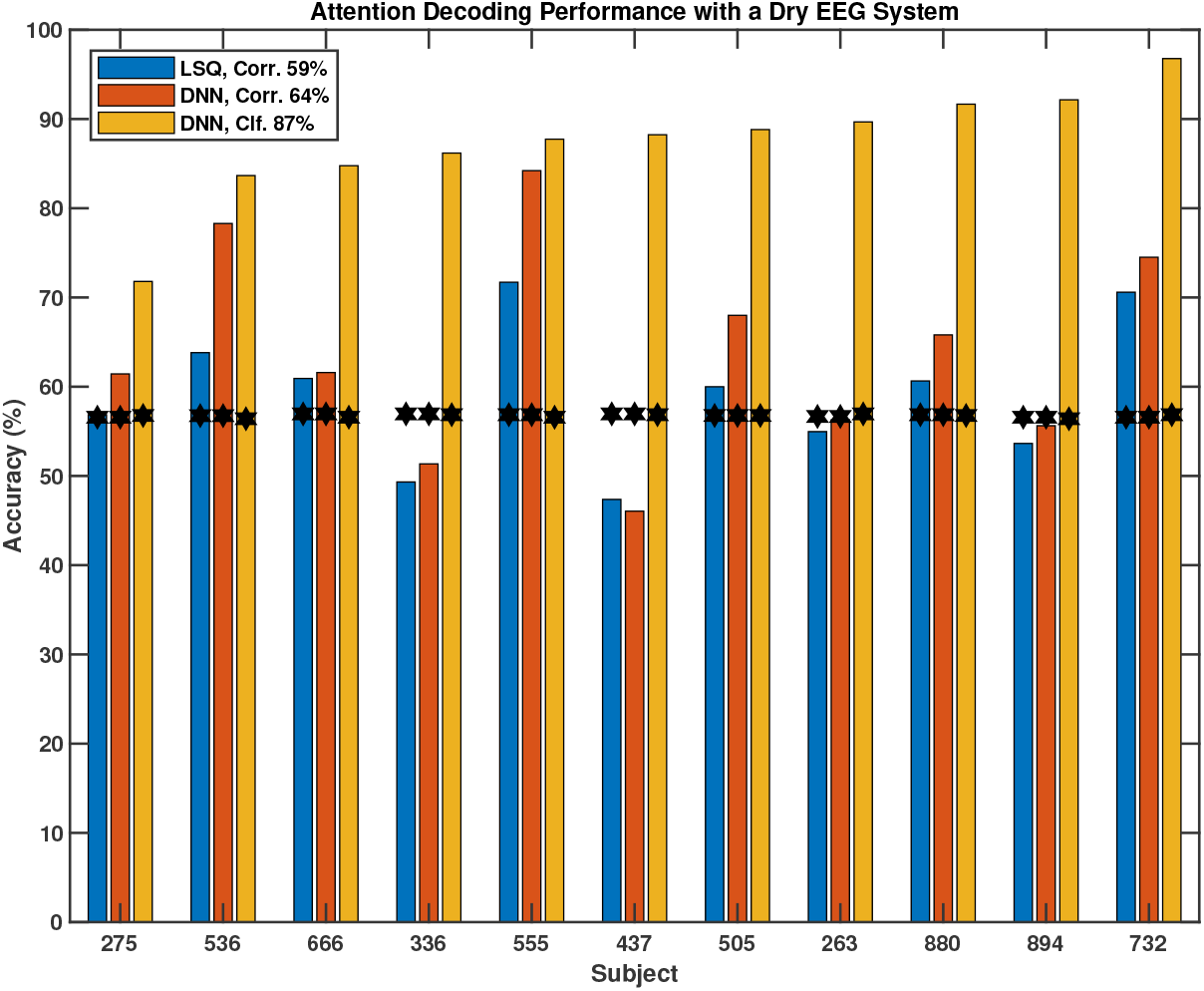
Per-subject attention-decoding accuracy using a dry EEG system. 10-second evaluation window, three algorithms: linear stimulus reconstruction (LSQ Corr.), non-linear stimulus reconstruction (DNN Corr.), and DNN classification (DNN Clf.). Chance performance is indicated by the black stars.

**Table 1.** Mean decoding accuracy for the three architectures and two EEG types. Standard deviations are shown in parentheses.

|  | Stimulus Reconstruction |  | Classifier |
| --- | --- | --- | --- |
|  | Linear | Nonlinear (DNN) | Nonlinear (DNN) |
| Wet EEG | 65% (10.3%) | 65% (9.7%) | 80% (5.5%) |
| Dry EEG | 59% (7.7%) | 64% (11.6%) | 87% (6.3%) |

### 3.2 Relationship between LSQ Regularization and Subject Decoding Accuracy

Consistent with [21], we found that regularization positively impacted LSQ decoding accuracy. While the decoding accuracies shown above were chosen from a set of three regularization values by examining one subject, post-hoc, we evaluated subject-level regularization values across a much larger range of candidate values. Fig 7 contains the median regularization values chosen by the internal cross validation loop plotted as a function of the decoding accuracy of the subject as determined using the optimal regularization value from the limited set of three values. There is a negative relationship between the subject’s median regularization parameter and the part accuracy achieved by the subject that is preserved across wet and dry modalities (*p* = 0.04,*p* = 0.13, respectively). Subjects with high performance required less weight penalization in the TRF construction process.

**Fig 7.**
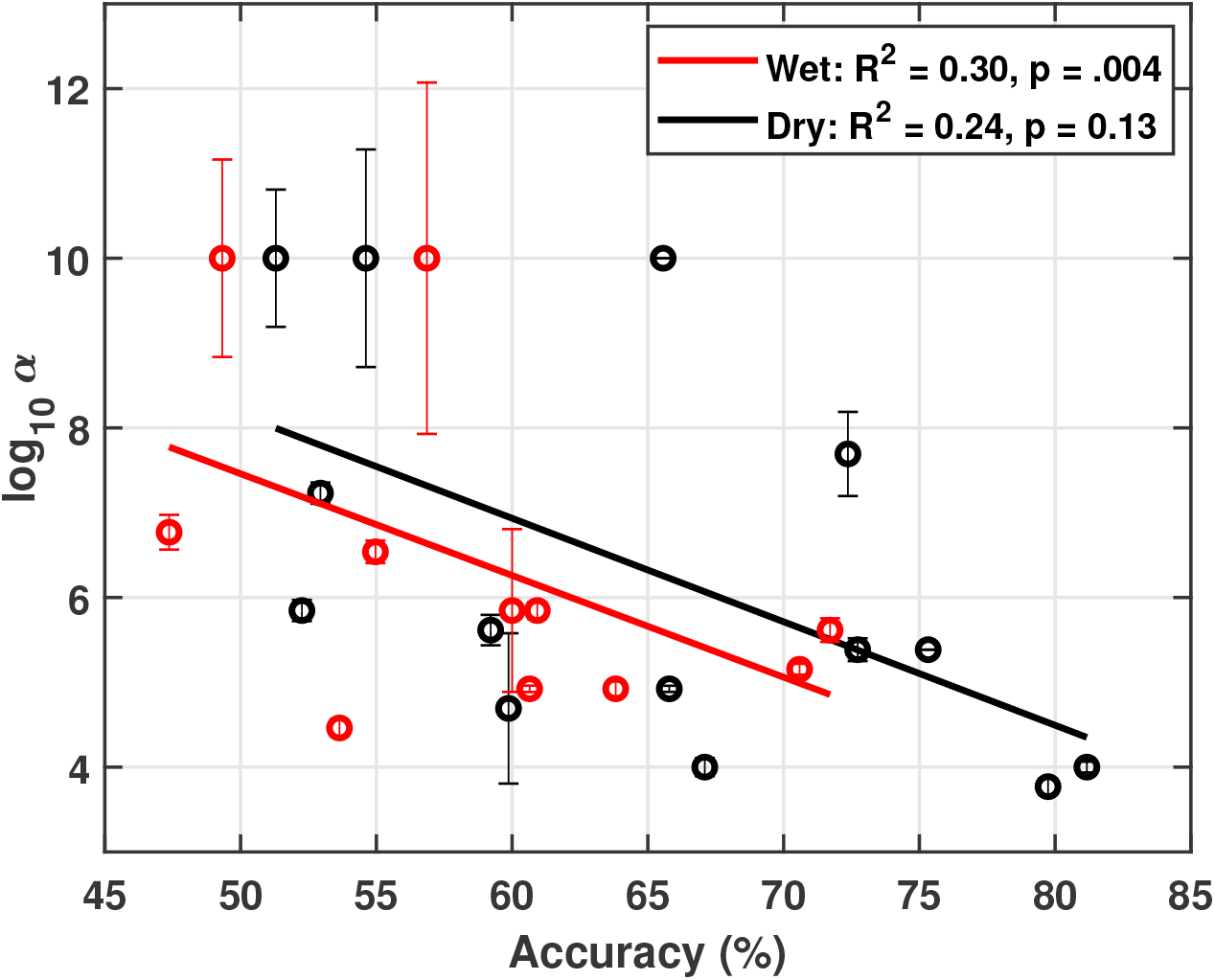
Median LSQ regularization parameters. Regularization parameter was taken across folds for both wet and dry EEG modalities, as a function of decoding performance.

### 3.3 Visualization of LSQ TRF

Through the larger sweep in regularization parameter described above we found that subjects with a higher tended to have smoother TRFs. Although performance on the test data drops as a result of increasing beyond the value used in the results section, the spatial progression of TRF intensity across time is more easily interpreted. For visualization, TRFs were constructed using a fixed regularization parameter of 10^9^ using LSQ estimation. The kernel length was expanded from 26 to 51 samples in order to ensure capturing the full temporal evolution of the transform, but on average only the first half of the TRFs showed substantial non-zero activity. As shown in Fig 8 as a 2D image and in Fig 9 as a series of headmaps, a TRF peak occurs at 200ms in the center of the head and dissipates afterwards. This timing is consistent with that reported in [13, 14, 20] where peaks near 200 ms also are shown.

**Fig 8.**
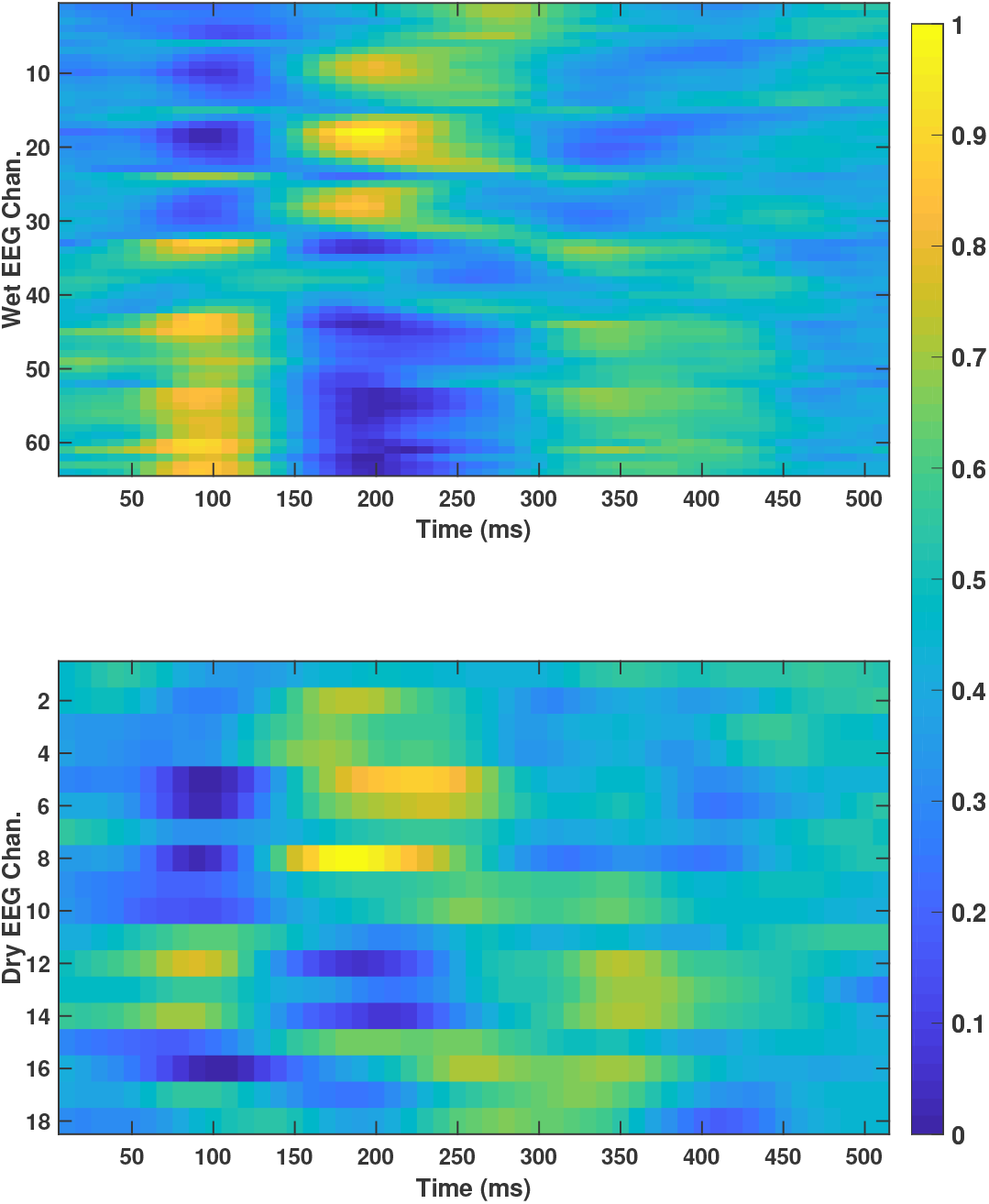
Grand average values of the LSQ TRFs across subjects. Top: 64-channel wet EEG; Bottom: 18-channel dry EEG.

**Fig 9.**
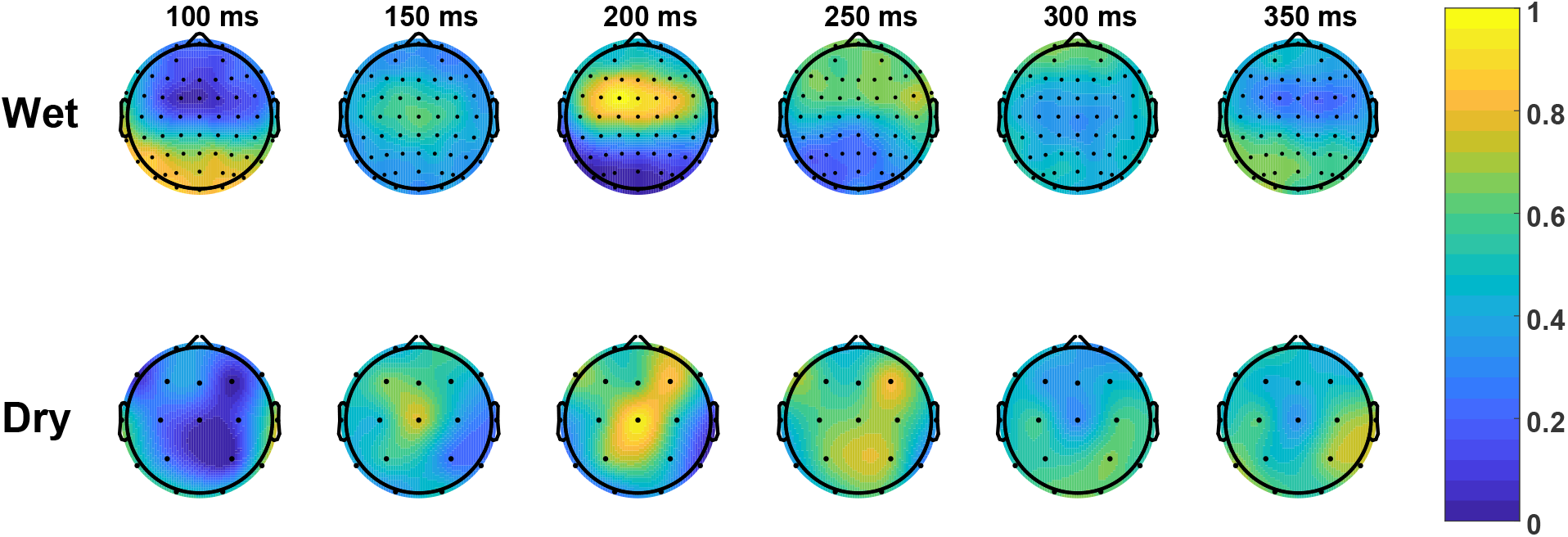
Grand average headmaps of the LSQ TRF values across subjects.

### 3.4 Channel Importance in the Convolutional DNN

While obtaining insight into why a DNN performs as it does remains a challenging research question, we can gain some understanding of the convolutional DNN by examining the filter weights of the first convolutional layer. Essentially, this convolution is creating a set of linear combinations of the input EEG and audio channels. The full convolutional weight matrix is 3-dimensional (kernel by input channel by output channel), but we can collapse the 3D matrix into one dimension in order to visualize it. First, we select the middle element of the three-point temporal kernel, and then take the absolute value of the weights. Next, we sum the convolutional weights along the input channel. Taking the wet EEG as an example, there are 64 EEG channels and an audio channel as the input and 64 channels as the output from the first convolutional layer. We renormalize the 64 EEG weights of the 65-element vector so the minimum weight is 0 and the maximum weight is 1 and then apply that normalization to the 65^*th*^ audio element. We compute the normalization separately for the wet and dry systems and per subject. Then, we average across the subjects and re-normalize again to a 0–1 range.

Fig 10 shows the mean absolute weights assigned to the wet and dry EEG datasets visualized as a headmap, with the weight for the audio channel indicated by the colored boxes below. Activated regions show some similarity to the LSQ TRF values in Fig 9. Specifically, for the wet-EEG case, the central peak for the DNN headmap is roughly co-located with the 200 ms peak for the LSQ TRF. For the dry-EEG case, the elongated activation area to the right of the mid-sagittal plane resembles that for the 250 ms LSQ TRF (although the central peak at 200 ms is not evident in the DNN weights). Since the DNN classifier takes both audio (envelope) and EEG as an input, the audio channel should be weighted highly, and we see this is the case with the wet electrode system yielding an audio weight of 1.0 and the dry electrode system yielding an audio weight of 0.95. This indicates that the network is utilizing both EEG and audio signals to make a decision. The electrodes with the highest weights were M1 and T8 for the wet and dry EEG systems, respectively.

**Fig 10.**
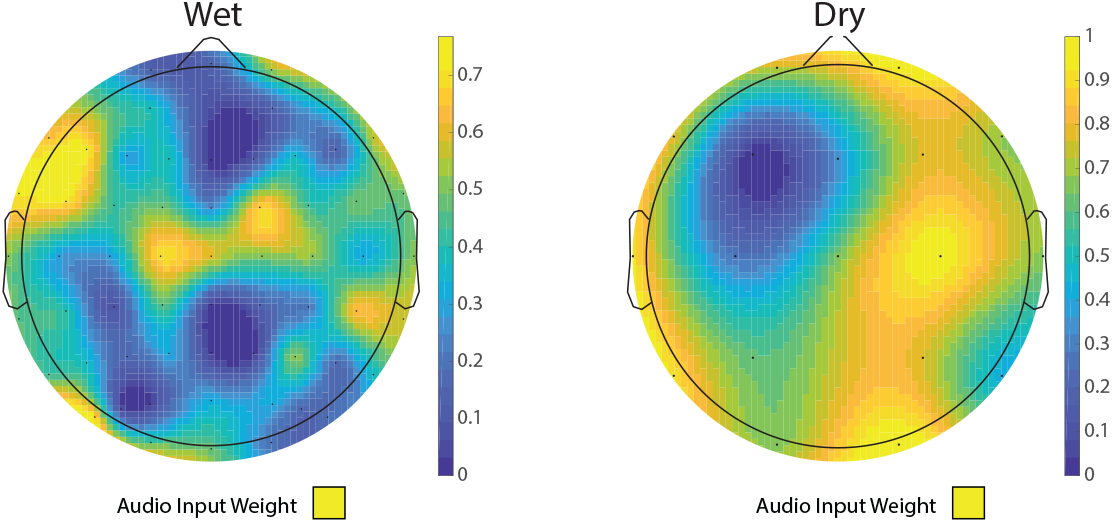
Grand average headmaps of the normalized mean convolutional weights for the wet and dry EEG systems for the DNN classifier network. The audio channel weights (bottom), also an input to the network, were 1.0 and 0.95 for the wet and dry systems, respectively.

## 4 Discussion

As shown in Figs 5 and 6, and Table 1, both the nonlinear and linear approaches yielded comparable performance under the stimulus-reconstruction architecture. Decoding accuracy in our study varied more with the subject than with the choice of these algorithms. Typically, either both approaches performed well on a subject (*e.g*., Subj. 555), or both performed poorly (*e.g*., Subj. 437). The DNN classifier approach dramatically outperformed the traditional segregated architecture in decoding accuracy (80% wet, 87% dry) with a performance advantage in all of the dry EEG cases and all but one of the wet EEG cases, and shows a smaller variance among the subjects. While the exact reason for this is unclear, future work includes further analysis of the DNN’s weights to better understand its learned similarity metric. In addition, comparison of the DNN classifier to a logistic-regression classifier could yield insight into the importance of non-linearities in the decoding process.

In regard to the two EEG systems, overall decoding performance is comparable between the wet electrode and dry electrode systems. This result is somewhat surprising given that the wet system contains more than three times as many channels (64 vs. 18), although earlier work has shown a channel reduction from 96 to 20 had limited effect on decoding accuracy [14]. Given these results, and recent studies that suggest that wet and dry EEG systems can deliver similar signal qualities (albeit with different systems than we used) [36], a practical integration of AAD into an unobtrusive, wearable hearing device seems to be an achievable, long-term goal.

Of the three approaches we considered, two explicitly involve a backward model, *i.e*., stimulus reconstruction. We did not test the forward decoding architecture in this paper for both empirical and theoretical reasons. In regard to the former, the forward decoding approach has shown slightly worse performance than the backward decoding approach [21]. Theoretically, this performance loss is understandable because the auditory stimulus is just one of many internal and external factors, none of which is known other than the audio, that influence the corresponding EEG waveform. By contrast, because the neural activity represented in the EEG data is at least in part due to an auditory stimulus, it is reasonable to filter out the non-auditory components but retain the auditory component. As an extreme example, assume a model for the transform from audio to a specific EEG channel as the envelope of the audio plus additive noise, with the noise independent at each lead. In this case, the forward problem requires predicting noise, whereas the backward problem allows averaging out the noise across all the leads to recover the auditory envelope.

The performance of the linear approach in our study was lower than that reported in previous studies, potentially due to differences in the experimental design and decoding parameters. One significant difference between the results reported here and in other publications is that our talkers were co-located, *i.e*., combined digitally and delivered from a single loudspeaker in front of the subject. Reduced spatial separation (down to 10°) has been shown to have a detrimental effect on decoding accuracy in low (−1.1, –4.1, and –7.1 dB) but not high (20 dB) SNR conditions [19], so it is not clear how strong an effect co-location had in this work. Other studies have included talkers at ±90° azimuth [13, 15–17, 19], ±60° [20, 21], ±30° [14], or ±10° [19]. We chose to use co-located talkers because this would provide a lower bound on decoding accuracy (from a spatial perspective) without extrapolating from an arbitrary separation angle.

A second potential reason for our relatively low linear decoding accuracy is that our correlation window (trial size) of 10 s and kernel length of 250 ms are shorter than those in some other experiments. Decoding accuracy previously has been shown to deteriorate with shortening trial sizes [17, 21, 35], and one-minute [13, 14] and 30-second [16, 19] windows are more common in the literature. Our choice of 10s was motivated by the fact that, a smaller window, eventually coupled with temporal smoothing such as that described in [29], will be necessary for use with a practical, low-latency AAD system. Least-squares kernels ranging from 250 ms [13, 17] to 500 ms [20, 21] have been reported, although no length has been shown to be optimal. We chose a 250 ms kernel based on early pilot data that did not indicate a significant improvement with an increase to 500 ms. Table 2 contains mean decoding accuracies for different correlation windows and kernel lengths to facilitate comparison to other AAD studies. Some improvement is seen with an increase in the correlation window length, but as with our pilot data, the kernel length had a negligible effect on performance.

**Table 2.** Mean decoding accuracy for the linear least-squares architecture with variations in the correlation window (10 s, 30 s) and the kernel size (250 ms, 500 ms). Standard deviations are shown in parentheses.

| Kernel Size |  | Correlation Window |  |
| --- | --- | --- | --- |
|  |  | 10 s | 30 s |
|  | 250 ms | Wet EEG: 65% (10.3%)<br>Dry EEG: 59% (7.7%) | Wet EEG: 72% (12.5%)<br>Dry EEG: 65% (14.4%) |
|  | 500 ms | Wet EEG: 66% (8.5%)<br>Dry EEG: 56% (8.1%) | Wet EEG: 70% (13.6%)<br>Dry EEG: 62% (13.1%) |

There are still several considerations in translating the decoding performance we are achieving to clinical utility. First, consistent with many other studies in the literature ([37] is an exception), we focused on normal hearing listeners and only included two hearing-impaired (HI) subjects. Interestingly, one of the HI subjects (with mild impairment) was in the top third of our cohort in terms of algorithm performance, while the other was in the bottom third. We will need to recruit a substantial group of HI subjects to evaluate these algorithms for their use. Second, there is significant variance in decoding performance across individuals. In our study, participants were randomly prompted to repeat the last sentence from the attended talker, but the recall accuracy was consistently high and does not explain the variation in performance. In addition to traditional hearing loss, other potential factors that could affect AAD performance include cochlear synaptopathy, cognitive ability (*e.g*., working memory), and fatigue. Such factors have been considered in the context of the variability of traditional hearing-aid performance/acceptance [38] and should be explored further in the context of AAD.

## 5 Conclusions

In conclusion, we have compared two different auditory decision architectures, one which employs a Pearson based similarity metric to compare the reconstructed stimulus with actual stimuli (using a linear or DNN-based reconstruction approach), and a second, novel version in which the similarity transform is learned as part of the optimization process in a convolutional neural network. Furthermore, we evaluated all three algorithms with both a wet and dry electrode EEG system using a two-talker AAD protocol. We found that the integrated decision-making architecture using a convolutional neural network yielded results comparable to state-of-the-art performance reported, and we have shown we can achieve this performance with both a wet and dry system where the talkers are not spatially separated. Future work includes evaluation of neural network architectures with around-the-ear [24] and in-ear [25] EEG electrodes. We also plan to employ transfer learning of network knowledge across subjects, and consider end-to-end neural network based architectures that combine both speaker separation and attention decoding, simply outputting the attended audio stream directly. This approach could be performed with single or multi-channel audio.

We plan to release both EEG datasets with baseline algorithms and benchmark performance metrics. We look forward to other research groups contributing their own analyses of this data in order to increase both the accuracy of decoding and shorten the latency of decoding. Improvements in both areas are needed for AAD to fulfill its promise as part of a complete, hearing-assistive system.

## Acknowledgment

DISTRIBUTION STATEMENT A. Approved for public release. Distribution is unlimited. This material is based upon work supported by the Assistant Secretary of Defense for Research and Engineering under Air Force Contract No. FA8702–15-D-0001. Any opinions, findings, conclusions or recommendations expressed in this material are those of the author(s) and do not necessarily reflect the views of the Assistant Secretary of Defense for Research and Engineering.

## References

1. Wilson BS, Tucci DL, Merson MH, O’Donoghue GM. Global hearing health care: New findings and perspectives. The Lancet. 2017;390(10111):2503–2515.

2. USVA. Annual Benefits Report Fiscal Year 2017; 2017. U.S. Department of Veterans Affairs, Veterans Benefits Administration.

3. Kochkin S. Customer satisfaction with hearing instruments in the digital age. The Hearing Journal. 2005;58(9):30–43.

4. Abrams H, Kihm J. An introduction to MarkeTrak IX: A new baseline for the hearing aid market. Hearing Review. 2015;22(6):16.

5. Lesica NA. Why Do Hearing Aids Fail to Restore Normal Auditory Perception? Trends in Neurosciences. 2018;.

6. Arlinger S. Negative consequences of uncorrected hearing loss – A review. International Journal of Audiology. 2003;42:2S17–2S20.

7. Mener DJ, Betz J, Genther DJ, Chen D, Lin FR. Hearing loss and depression in older adults. Journal of the American Geriatrics Society. 2013;61(9):1627–1629.

8. Andrade CC, Pereira CR, DA SILVA PA. The silent impact of hearing loss: Using longitudinal data to explore the effects on depression and social activity restriction among older people. Ageing & Society. 2017; p. 1–22.

9. Aiken SJ, Picton TW. Human cortical responses to the speech envelope. Ear and Hearing. 2008;29(2):139–157.

10. Ding N, Simon JZ. Neural coding of continuous speech in auditory cortex during monaural and dichotic listening. Journal of Neurophysiology. 2012;107(1):78–89.

11. Golumbic EMZ, Ding N, Bickel S, Lakatos P, Schevon CA, McKhann GM, et al. Mechanisms underlying selective neuronal tracking of attended speech at a “cocktail party”. Neuron. 2013;77(5):980–991.

12. Mesgarani N, Chang EF. Selective cortical representation of attended speaker in multi-talker speech perception. Nature. 2012;485(7397):233.

13. O’Sullivan JA, Power AJ, Mesgarani N, Rajaram S, Foxe JJ, Shinn-Cunningham BG, et al. Attentional selection in a cocktail party environment can be decoded from single-trial EEG. Cerebral Cortex. 2015;25(7):1697–1706.

14. Mirkovic B, Debener S, Jaeger M, De Vos M. Decoding the attended speech stream with multi-channel EEG: Implications for online, daily-life applications. Journal of Neural Engineering. 2015;12(4):046007.

15. Aroudi A, Mirkovic B, De Vos M, Doclo S. Auditory attention decoding with EEG recordings using noisy acoustic reference signals. In: Acoustics, Speech and Signal Processing (ICASSP), 2016 IEEE International Conference on. IEEE; 2016. p. 694–698.

16. Van Eyndhoven S, Francart T, Bertrand A. EEG-informed attended speaker extraction from recorded speech mixtures with application in neuro-steered hearing prostheses. IEEE Transactions on Biomedical Engineering. 2017;64(5):1045–1056.

17. Biesmans W, Das N, Francart T, Bertrand A. Auditory-inspired speech envelope extraction methods for improved EEG-based auditory attention detection in a cocktail party scenario. IEEE Transactions on Neural Systems and Rehabilitation Engineering. 2017;25(5):402–412.

18. Mendoza CF, Segar A. Decoding Auditory Attention from Multivariate Neural Data using Cepstral Analysis. Lund University, Dept. of Mathematical Statisics; 2018.

19. Das N, Bertrand A, Francart T. EEG-based auditory attention detection: Boundary conditions for background noise and speaker positions. Journal of Neural Engineering. 2018;.

20. Fuglsang SA, Dau T, Hjortkjær J. Noise-robust cortical tracking of attended speech in real-world acoustic scenes. NeuroImage. 2017;156:435–444.

21. Wong DD, Fuglsang SA, Hjortkjær J, Ceolini E, Slaney M, De Cheveigne A. A comparison of regularization methods in forward and backward models for auditory attention decoding. Frontiers in Neuroscience. 2018;12:531.

22. Akram S, Presacco A, Simon JZ, Shamma SA, Babadi B. Robust decoding of selective auditory attention from MEG in a competing-speaker environment via state-space modeling. NeuroImage. 2016;124:906–917.

23. O’Sullivan J, Chen Z, Sheth SA, McKhann G, Mehta AD, Mesgarani N. Neural decoding of attentional selection in multi-speaker environments without access to separated sources. In: Engineering in Medicine and Biology Society (EMBC), 2017 39th Annual International Conference of the IEEE. IEEE; 2017. p. 1644–1647.

24. Bleichner MG, Mirkovic B, Debener S. Identifying auditory attention with ear-EEG: cEEGrid versus high-density cap-EEG comparison. Journal of Neural Engineering. 2016;13(6):066004.

25. Fiedler L, Wöstmann M, Graversen C, Brandmeyer A, Lunner T, Obleser J. Single-channel in-ear-EEG detects the focus of auditory attention to concurrent tone streams and mixed speech. Journal of Neural Engineering. 2017;14(3):036020.

26. Baltzell LS, Horton C, Shen Y, Richards VM, D’Zmura M, Srinivasan R. Attention selectively modulates cortical entrainment in different regions of the speech spectrum. Brain Research. 2016;1644:203–212.

27. de Taillez T, Kollmeier B, Meyer BT. Machine learning for decoding listeners’ attention from electroencephalography evoked by continuous speech. European Journal of Neuroscience. 2017;.

28. de Cheveigné A, Wong DD, Di Liberto GM, Hjortkjær J, Slaney M, Lalor E. Decoding the auditory brain with canonical component analysis. NeuroImage. 2018;172:206–216.

29. Miran S, Akram S, Sheikhattar A, Simon JZ, Zhang T, Babadi B. Real-time tracking of selective auditory attention from M/EEG: A bayesian filtering approach. Frontiers in Neuroscience. 2018;12.

30. Horwitz-Martin RL, Quatieri TF, Godoy E, Williamson JR. A vocal modulation model with application to predicting depression severity. In: Wearable and Implantable Body Sensor Networks (BSN), 2016 IEEE 13th International Conference on; 2016. p. 247–253.

31. Pedregosa F, Varoquaux G, Gramfort A, Michel V, Thirion B, Grisel O, et al. Scikit-learn: Machine Learning in Python. Journal of Machine Learning Research. 2011;12:2825–2830.

32. Ioffe S, Szegedy C. Batch normalization: Accelerating deep network training by reducing internal covariate shift. arXiv preprint arXiv:150203167. 2015;.

33. Srivastava N, Hinton G, Krizhevsky A, Sutskever I, Salakhutdinov R. Dropout: A simple way to prevent neural networks from overfitting. The Journal of Machine Learning Research. 2014;15(1):1929–1958.

34. Clevert DA, Unterthiner T, Hochreiter S. Fast and accurate deep network learning by exponential linear units (elus). arXiv preprint arXiv:151107289. 2015;.

35. Zink R, Proesmans S, Bertrand A, Van Huffel S, De Vos M. Online detection of auditory attention with mobile EEG: Closing the loop with neurofeedback. bioRxiv. 2017; p. 218727.

36. Kam JW, Griffin S, Shen A, Patel S, Hinrichs H, Heinze HJ, et al. Systematic comparison between a wireless EEG system with dry electrodes and a wired EEG system with wet electrodes. NeuroImage. 2018;.

37. Dau T, Maercher Roersted J, Fuglsang S, Hjortkjær J. Towards cognitive control of hearing instruments using EEG measures of selective attention. The Journal of the Acoustical Society of America. 2018;143(3):1744.

38. Tremblay K, Miller C. How neuroscience relates to hearing aid amplification. International Journal of Otolaryngology. 2014;2014.

